# Hemodynamic changes in response to excitatory and inhibitory modulations by transcranial magnetic stimulation at the human sensorimotor cortex

**DOI:** 10.1101/2020.12.19.423591

**Authors:** Hsin-Ju Lee, Mikko Nyrhinen, Risto J. Ilmoniemi, Fa-Hsuan Lin

**Affiliations:** Physical Sciences Platform, Sunnybrook Research Institute, Toronto, ON, Canada; Department of Medical Biophysics, University of Toronto, Toronto, ON, Canada; Department of Neuroscience and Biomedical Engineering, Aalto University, Espoo, Finland

**Author notes:** Correspondence: Hsin-Ju Lee, Ph.D., Address: Rm. S605, 2075 Bayview Avenue, Toronto, ON, M4N 3M5, Canada.

## Abstract

Transcranial magnetic stimulation (TMS) can non-invasively induce both excitatory and inhibitory neuronal activity. However, the neurophysiological basis for both kinds of modulation remains elusive. In this study, with a controlled dosage over the 30-s interval, we elicited excitatory and inhibitory TMS modulations over the human primary motor cortex (M1) with TMS bursts of high (10-Hz and 30-Hz) and low frequency (0.5-Hz), respectively, and took functional magnetic resonance images (fMRI). Excitatory and inhibitory modulations were evidenced by changes in motor evoked potentials (MEP). Significantly increased fMRI signal at M1 was only detected under excitatory high-frequency TMS but not during inhibitory low-frequency TMS. The supplementary motor area (SMA) had significant fMRI signal changes after both excitatory and inhibitory TMS. The topology of the activated M1 and SMA matched the activated sensorimotor network during voluntary movement. The precuneus was selectively activated with bursts of five TMS pulses. These findings demonstrated the asymmetric hemodynamic responses to excitatory and inhibitory TMS modulations with region-dependent relationships between the local fMRI signal changes and TMS dosage over different time scales.

## Introduction

Excitatory and inhibitory responses are two opposite outcomes of neuronal modulation. Both responses can be manifested in behaviors, which are likely supported by different orchestrated neuronal activity across regions. Transcranial magnetic stimulation (TMS) can non-invasively excite or inhibit activity at a focal brain area by delivering brief magnetic pulses [1]. Specifically, delivering TMS pulses over the primary motor cortex (M1) at a slow rate (≤ 1 Hz) can cause an inhibitory effect on the motor evoked potentials (MEP) by attenuating its amplitude [2, 3]. Similarly, delivering TMS pulses at a fast rate (≥ 5 Hz) excites the targeted cortical area to cause the MEP amplitude increase [4]. How brain activity causes MEP changes in TMS excitatory and inhibitory modulations remains elusive.

The concurrent recording of TMS with electroencephalography (EEG) [3, 5] and functional magnetic resonance imaging (fMRI) [6, 7] can measure the brain electrophysiological and hemodynamic responses to TMS modulations, respectively. For example, Casula et al. (2014) reported a significant decrease in the MEP amplitude with a significant increase in the EEG evoked responses following 1,200 pulses of 1-Hz repetitive TMS over the M1. In concurrent TMS–fMRI recording, a nonlinear relationship between TMS stimulation strength and fMRI signal strength was found by delivering single TMS pulses to M1. Significantly increased fMRI signal at the sensorimotor network was only observed when the TMS stimulation strength exceeded the resting motor threshold (rMT). In contrast, a trend of reduced fMRI signal associated with sub-threshold TMS stimulation was observed [7]. These findings suggest different neurophysiological responses to excitatory and inhibitory neuromodulations.

Previous TMS-fMRI studies have revealed that the dependency of the brain responses on the TMS intensity [6–10]. Specifically, TMS over the M1 induces cortical and subcortical fMRI signal changes [7, 9, 10]. Interestingly, the TMS target site shows a positive fMRI signal change only when it is stimulated at a relatively high TMS intensity. These observations reflect the complex interaction between the TMS stimulation dosage, the functional state of the targeted brain area, and neurovascular coupling characteristics.

Here, targeting a single brain area with a controlled TMS dosage over 30 seconds, we used concurrent TMS–fMRI to reveal brain responses to excitatory and inhibitory TMS modulations in the sensorimotor network. We chose fMRI to monitor brain activity because of its good spatial resolution. In contrast, while EEG suffers from uncertainty in localizing brain activity, it has a high temporal resolution and high sensitivity to neuronal activity. We first hypothesized that the spatial distribution of the sensorimotor network in response to TMS stimulation matches with the network in voluntary movements. We also hypothesized that the fMRI signal changes due to TMS modulations in the sensorimotor network were similar in magnitude and polarity. The former hypothesis was based on the similar sensorimotor network topology in the resting state [11] and during both voluntary [12] and involuntary movements [7, 13]. The latter hypothesis was based on the temporally correlated fMRI dynamics in the regions within the sensorimotor network [14]. Understanding how excitatory and inhibitory neuromodulations corresponds to fMRI signal changes can help decipher the neuronal mechanism underlying fMRI responses [15].

## Methods

### Participants

Twelve healthy participants (5 females; age: 30 ± 5.8 years) participated in the experiment after giving written informed consent. None of the participants reported a history of neuropsychiatric disease or clinical evidence of neurological dysfunction. One of the participants was left-handed. This study followed the Declaration of Helsinki guidelines and was approved by the ethics committee of Aalto University.

### Experimental design

The experiment consisted of two sessions. In the TMS–fMRI session, fMRI was acquired using a customized 8-channel head coil array compatible with TMS [16]. In the second session, fMRI without TMS was acquired using a commercial 32-channel head coil array (Siemens, Erlangen, Germany).

The first session comprised six runs of fMRI with TMS. A *T*_1_-weighted anatomical scan and a field map scan were acquired. The TMS runs (**Figure 1a**) started with a 10-s interval of “off” (without TMS delivery) followed by five alternating 30-s “on” (with TMS delivery) and 30-s “off” blocks. In each TMS block, 15 pulses in total were delivered in order to control the TMS dose. Five conditions of TMS pulse delivery were administered: “0.5 Hz”, where TMS pulses were equally separated by 2 s; “10 Hz, 3 pulses per burst (ppb)”, where TMS pulses were delivered in five bursts with three pulses with 0.1-s inter-pulse interval (ipi) and 6-s inter-burst interval (ibi); “10-Hz–5-ppb”, where TMS pulses were delivered in three bursts with five pulses with 0.1-s ipi and 10-s ibi; “30-Hz–3-ppb”, where TMS pulses were delivered in five bursts with three pulses with 0.033-s ipi and 6-s ibi; “30-Hz–5-ppb”, where TMS pulses were delivered in three bursts with five pulses with 0.033-s ipi and 10-s ibi. Five TMS blocks were randomized and the stimulation intensity was 100% of the individual’s rMT. Within each TR (2 s), the acquisition of each MRI volume was completed between 0 and 1 s after the onset of each volumetric acquisition, whereas TMS pulses were delivered between 1 and 1.9 s. The last 0.1-s interval was set to avoid the TMS pulse interference to MRI acquisition based on previous TMS–fMRI studies [17]. **Figure 1b** shows the timing diagram of these TMS conditions. TMS pulse delivery was synchronized to the MRI trigger signal by using Presentation (version 20.0; Neurobehavioral systems, Albany, CA, USA). A total of 155 MRI volumes were acquired for each 310-s run. Participants were asked to stay still while resting and to fixate on a dot at the center of the visual field throughout the entire session.

**Figure 1.**
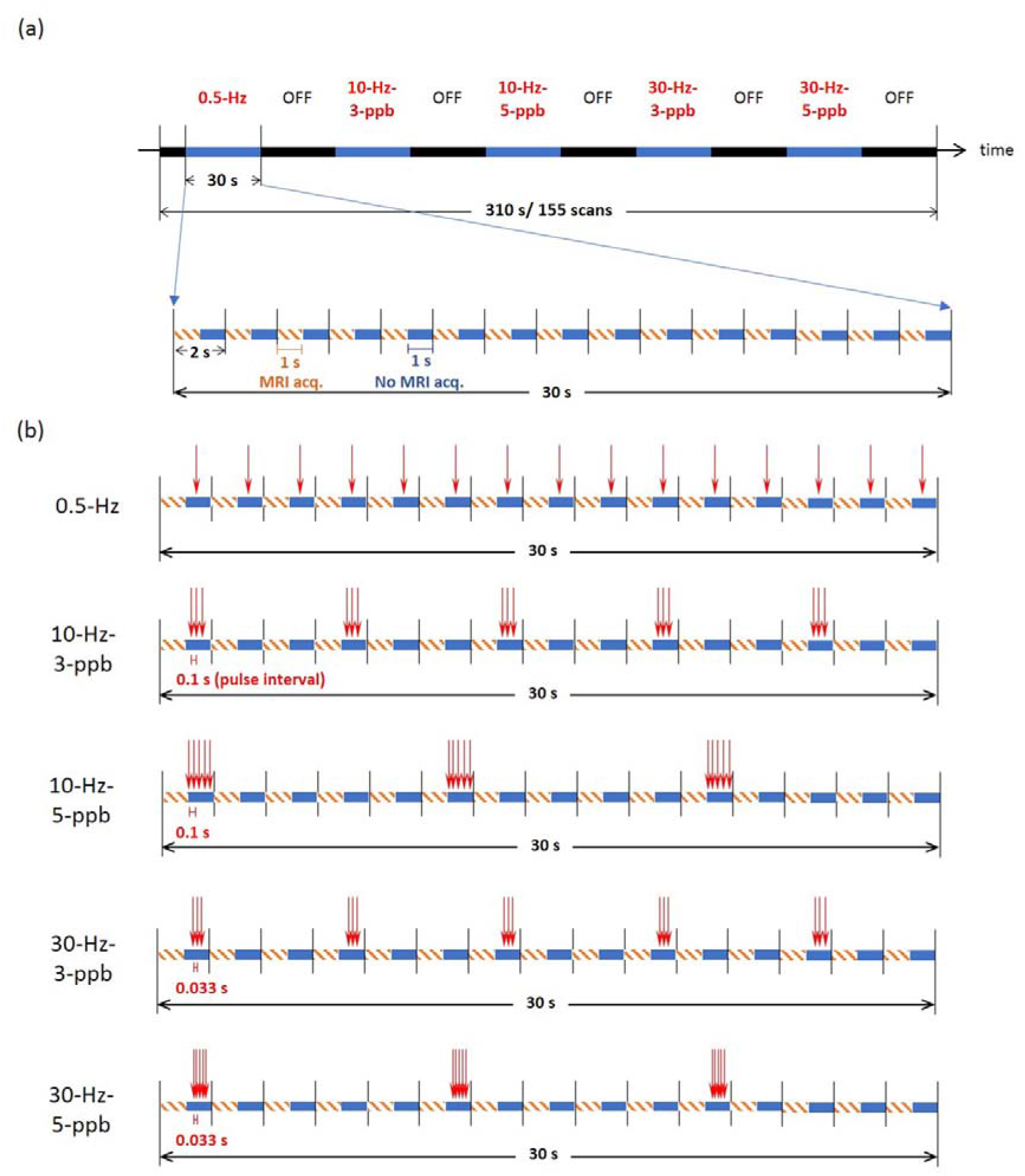
TMS pulse sequences with fMRI acquisitions. (a) Each fMRI run lasted 310 s and started with a 10-s interval of “off” (blacks segments represent intervals without TMS delivery) followed by five alternating 30-s TMS (blue segments) and 30-s “off” blocks. Five different conditions of “TMS” stimuli were administered in a random order. (b) Within each TR (2 s), the acquisition of each MRI volume was completed between 0 and 1 s after the onset of each volumetric acquisition (orange dashed line), whereas TMS pulses were delivered between 1 and 1.9 s during the MRI quiet period (blue). In each TMS block, a total of 15 pulses were delivered.

In the second session, participants were instructed to tap their index finger to thumb bilaterally at a self-paced speed (four 30-s blocks separated by 30 s). Two runs of EPI were acquired to functionally define the region of interest. Additionally, an anatomical scan was collected.

### Coil array construction and MRI acquisition

All data were acquired on a 3T MRI system (Skyra, Siemens Healthcare, Erlangen, Germany). We developed a dedicated 8-channel surface coil array [16] integrated with an MRI-compatible TMS coil (MRi-B91, MagVenture, Denmark) connected to a TMS stimulator system (MagPro X100, MagVenture, Denmark). In the first session, intermittent echo-planar imaging (TR = 2 s, TE = 26 ms, flip angle = 90°, 14 slices, 2-mm isotropic resolution) was prescribed to cover the left M1. The image planes were aligned in parallel with the TMS coil plane. Each MRI volume was completed between 0 and 1 s after the onset of each volumetric acquisition. In the second session, finger-tapping fMRI data was acquired with a single-shot EPI sequence (TR = 2000 ms, TE = 29 ms, flip angle = 30°, 33 slices, 3.3-mm slice thickness, 3.3×3.3-mm^2^ in-plane resolution). Structural images for each participant were acquired using a T1-weighted MPRAGE sequence (FOV: 256 x 256 mm2; TR = 2530 ms; TE = 3.3 ms; TI = 1100 ms; FA = 7°; BW = 200 Hz/pixel; 192 sagittal slice).

### TMS and electromyography recordings

Magnetic stimulation was performed with an MR-compatible TMS coil system. The stimulator generated biphasic pulses of approximately 280 μs in duration. The TMS coil was connected with an extension cable (length = 8 m) through a radiofrequency filter tube to a filter box mounted on the filter plate. This box also contained hardware to suppress leakage currents below 1 μA (Magventure, Farum, Denmark). The TMS coil was snapped on top of our developed MRI coil array and held steadily by a coil holder (MRi Coil Holder, MagVenture, Farum, Denmark). The coil plane was oriented to be tangential to the scalp and its normal direction was approximately 45° away from the midline between hemispheres. Each participant lay supine on a customized head holder with cushion. The head movement was restricted by foam-padded and vacuum cushions and the participant wore earplugs and earmuffs as hearing protection to reduce the perceived acoustic noise throughout the experiment.

The TMS target area was the cortical representation area of the right first dorsal interosseous (FDI) muscle on the M1. This area was identified for each participant by observing clear movements of the right index finger when the participant sat on the MRI bed outside the MRI bore. The location and orientation of the TMS coil when such movements were observed were marked on a hairnet supported by a net cap worn by the participant. Then, the participant lay on the MRI bed outside the MRI bore and we delivered TMS pulses to ensure clear movement of the participant’s right index finger. The motor threshold was determined for each participant when she/he lay inside the MRI scanner. The individual’s rMT was determined by varying the TMS pulse intensity until the electromyographic (EMG) MEP of the right FDI muscle was larger than 50 μV peak-to-peak in five out of ten stimuli [18].

EMG was recorded via one pair of radiotranslucent electrodes (EL508, Biopac Systems Inc., USA) with the active electrode on the FDI of the right index finger and the reference electrode which was about 2 cm from the active electrode. The ground electrode was placed on the participant’s wrist on top of the ulna bone. EMG signals were sampled at 2 kHz, amplified, filtered (1 to 150 Hz) and recorded by an electrocardiogram amplifier (MP-150 and ECG100C, Biopac Systems Inc., U.S.A.) and a computer for off-line analysis.

### MEP analysis

MEP data were analyzed by Matlab (The MathWorks, Inc., Natick, MA, USA). The peak-to-peak (in μV) amplitudes were calculated from the time interval between 20 and 40 ms after the TMS stimulus onset. Only MEP peak-to-peak amplitudes in the first pulse of each TMS burst were selected to calculate their mean and variance in each TMS condition.

### Functional MRI analysis

Image pre-processing was performed using SPM12 (Wellcome Department of Imaging Neuroscience, Institute of Neurology, London, UK), including slice-timing correction, motion correction, coregistration between functional and anatomical data, spatial normalization to the MNI space, and smoothing with a 6-mm full-width-half-maximum isotropic Gaussian kernel. The pre-processed datasets were then analyzed using the General Linear Model (GLM). A high-pass filter (1/128 Hz) was applied to remove low-frequency noise from the time series data, and serial correlations were corrected using an autoregressive AR(1) model. Based on the TMS conditions (TMS–fMRI experiment) or paced tapping blocks (finger-tapping experiment), a box-car function with time instants with TMS pulse delivery or finger movement were defined. This time series model was convolved with SPM’s canonical hemodynamic response function in the first-level analysis.

The contrast images of interest were entered into the second-level analyses for group results. A one sample *t*-test was used to assess brain activation in the finger tapping task. A repeated-measures ANOVA was administered for the TMS effects. *F*-maps constructed for each condition with a threshold of 0.005 were used to detect the significant fMRI signal changes evoked by TMS. Post-hoc *t*-maps were then used to determine brain activation and deactivation with a Bonferroni corrected threshold of 0.025. The reported statistical significance was corrected for multiple comparison by using a false discovery rate (FDR)-adjusted *p* < 0.05.

### Regions-of-interest analysis

Regions of interest (ROI) were selected on the basis of active regions in the fingertapping task at an FDR-corrected *p* < 0.05. As the coverage of the customized MR receive array was limited, only the fMRI signal intensities at the left SMC and SMA were extracted. The fMRI signals were extracted from each participant’s ROIs and averaged across all participants to study the effects of TMS.

### Statistical analysis

Statistical tests were performed using Statistical Package for Social Sciences (version 17.0; SPSS Inc., Chicago, IL, USA). We started the statistical analysis by testing the normality of data distribution with the Shapiro–Wilk’s test. Data were log-10-transformed if not distributed normally, as suggested by authors of previous TMS studies [19, 20]. The repeated-measures ANOVA test and *t*-test were used to compare the means when the variance across conditions was homogeneously ascertained by Levene’s test or an *F*-test. Otherwise, nonparametric Friedman test and Wilcoxon Signed Ranks test were used to compare the means.

## Results

TMS was delivered at 100% of the rMT with 0.5-Hz pulse rate to cause inhibitory modulation and with 10- and 30-Hz pulse rates to cause excitatory modulation. The pulses were delivered while the participants were at rest in the 3-T MRI scanner with concurrent fMRI scanning. Functional MRI signal changes due to TMS modulations were compared with the sensorimotor network subserving a voluntary finger-tapping task.

None of the participants reported adverse effects from the experimental procedure. The mean rMT was 77%, ranging from 59% to 99% of the maximum output of the TMS stimulator. Because of severe head motion, three datasets were excluded from further analysis. Nine participants were included in the group-level analysis. One participant was excluded from the MEP analysis because of a technical problem in EMG recording.

### Behavioral responses

Baseline MEP was taken for each participant inside the MRI before any TMS modulation. Significant MEPs were detected in the baseline and all TMS conditions (**Figure 2**). The MEP values were presented in the log-10 scale MEP distributions toward normality [19, 20]. We found no significant difference between MEPs caused by different frequencies (10-Hz vs. 30-Hz; two-way repeated measures ANOVA: *F_(1,7)_* = 4.077, *p* = 0.083) or the pulse numbers (5-ppb vs. 3-ppb; two-way repeated-measures ANOVA: *F*_(1,7)_ = 0.965, *p* = 0.359). There was no interaction between the TMS frequencies and pulse numbers (two-way repeated-measures ANOVA: *F_(1,7)_* = 0.814, *p* = 0.397). Therefore, we grouped responses evoked by 10-Hz and 30-Hz TMS pulses together as the response in the high-frequency (HF) condition. The 0.5-Hz TMS pulses were chosen as the low-frequency (LF) condition. MEPs at both HF and LF conditions were compared with the baseline MEP. Significant differences among the three conditions were found (one-way repeated-measures ANOVA: *F*_(2,14)_ = 12.542, *p* = 0.001). Post-hoc paired *t*-tests suggested that the average MEP amplitude in the HF condition was significantly larger than that in the baseline (one-tailed *t*-test: *t*_(7)_ = −2.008, *p* = 0.042). The average amplitude of MEP in the LF condition was significantly smaller than that in the baseline (one-tailed *t*-test: *t*_(7)_ = −2.727, *p* = 0.015) and that in the HF condition (one-tailed *t*-test: *t*_(7)_ = −6.170, *p* < 0.001). The MEP measures were in line with excitatory and inhibitory modulations by HF and LF TMS pulses, respectively.

**Figure 2.**
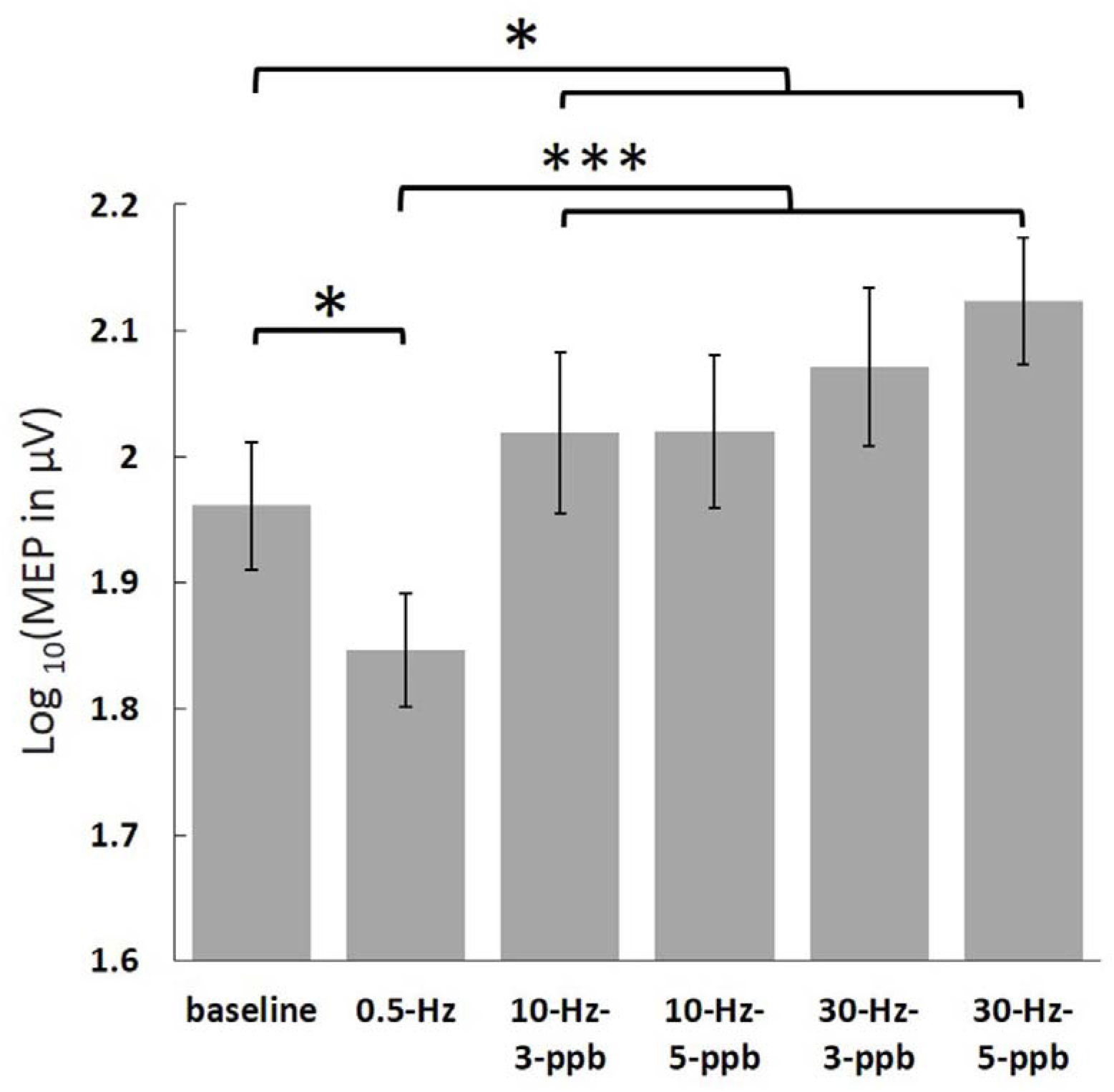
Log-transformed MEP amplitudes in the baseline and in the individual TMS conditions. Significant MEPs were detected in all conditions. The log-transformed MEP amplitude in the LF condition was significantly smaller than in the baseline and HF conditions. In contrast, the log-transformed MEP amplitude in the HF condition was significantly larger than in the baseline. Error bars denote standard errors of the mean (SEM). *: *p* < 0.05. ***: *p* < 0.001.

### Functional MRI signals elicited by TMS

Maps of brain activation and deactivation evoked by TMS are shown in **Figure 3a** and **Table 1**. For each TMS condition, significant fMRI signal changes were first identified using an undirected *F*-test. Then, post-hoc *t*-tests were used to examine the polarity of fMRI signal changes. Brain activation indicated by the increased fMRI signals were found at the sensorimotor cortex (SMC) ipsilateral to the TMS target locus in all 10-Hz and 30-Hz conditions, but not in the LF (0.5-Hz) condition. Compared to the fMRI signal without delivering any TMS, the fMRI signal in the supplementary motor area (SMA) was significantly increased in the 10-Hz–5-ppb, 30-Hz–3-ppb, and 30-Hz–5-ppb conditions. The significantly decreased fMRI signals were found at the left anterior middle frontal gyrus in the 10-Hz–5-ppb condition and in the medial wall of right SMC in the 30-Hz–3-ppb condition. In the HF condition (all 10- and 30-Hz conditions), significantly increased fMRI signals were found at the SMA and the left SMC. Significantly reduced fMRI signals were also found at bilateral anterior middle frontal gyrus, the bilateral superior parietal lobule, as well as the medial portion and the lateral surface of contralateral SMC in the HF condition (**Figure 3b** and **Table 1**).

**Figure 3.**
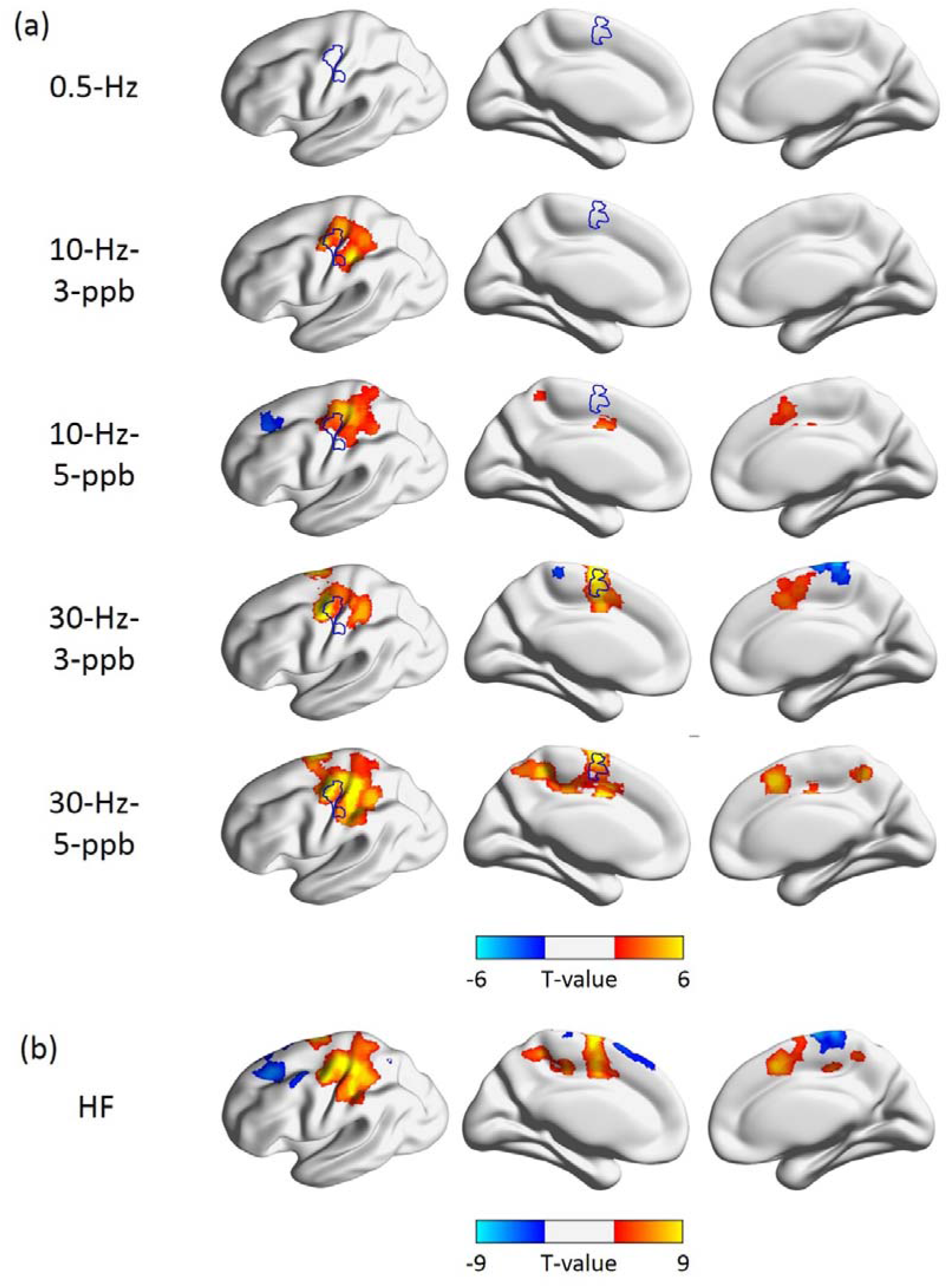
Distributions of significant fMRI signal changes in the individual TMS conditions (a) and the HF condition (b). (a) Positive fMRI signals were found at the SMC ipsilateral to the TMS target locus in all 10-Hz and 30-Hz conditions, but not in the 0.5-Hz condition. The fMRI signal in the SMA was significantly increased in the 10-Hz–5-ppb, 30-Hz–3-ppb, and 30-Hz–5-ppb conditions. The significantly negative fMRI signals were found at the left anterior middle frontal gyrus in the 10-Hz–5-ppb condition and in the medial wall of right SMC in the 30-Hz–3-ppb condition. (b) Significantly increased fMRI signal was found at the SMA and the left SMC in the HF condition. Contrarily, significantly reduced fMRI signals were found at bilateral anterior middle frontal gyrus, the bilateral superior parietal lobule, the medial portion and the lateral surface of contralateral SMC. The thresholds for these maps are false discovery rate (FDR)-adjusted *p* < 0.05. Blue contours indicate the activated brain areas in the finger-tapping task.

**Table 1.**
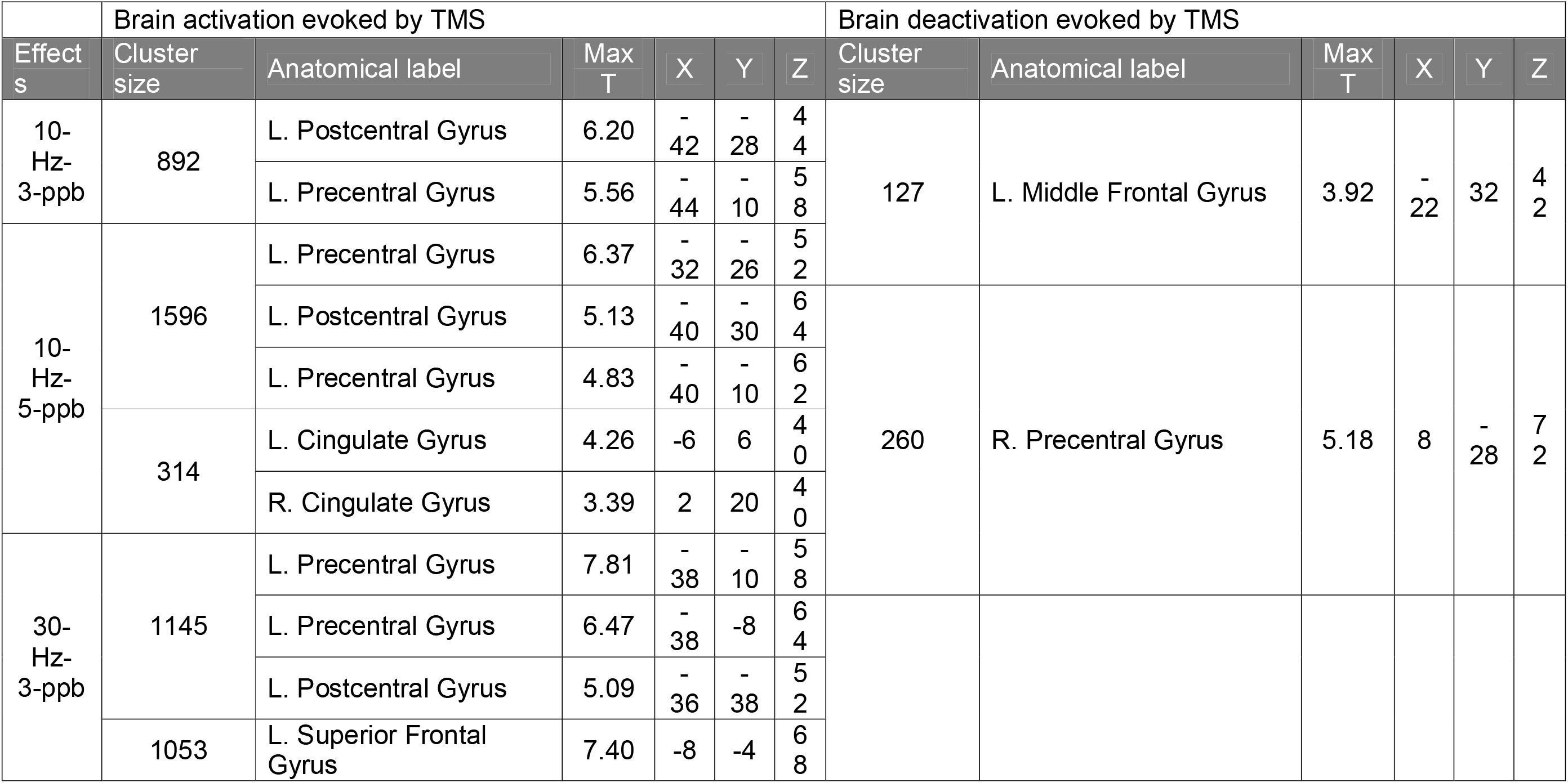
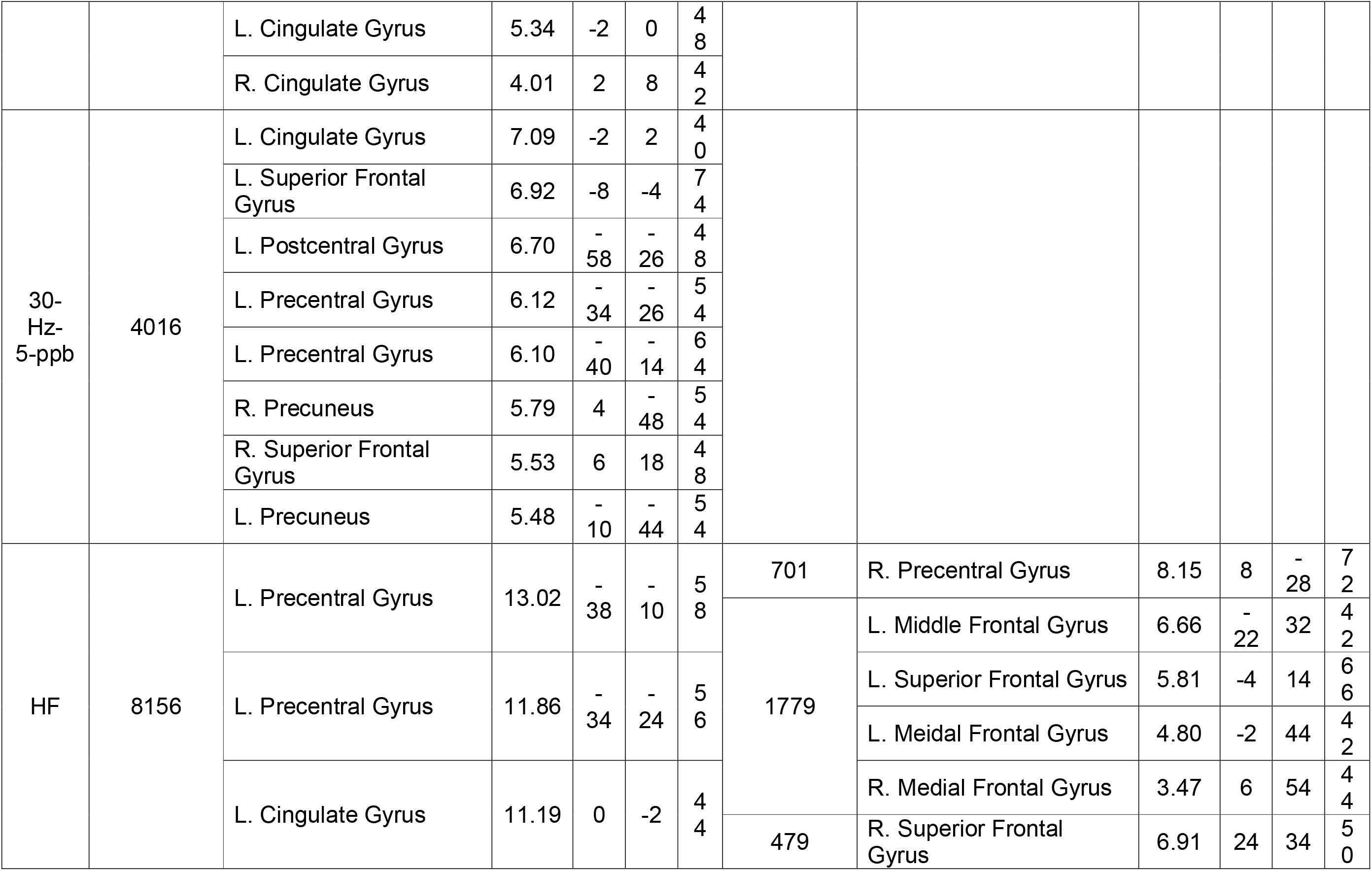
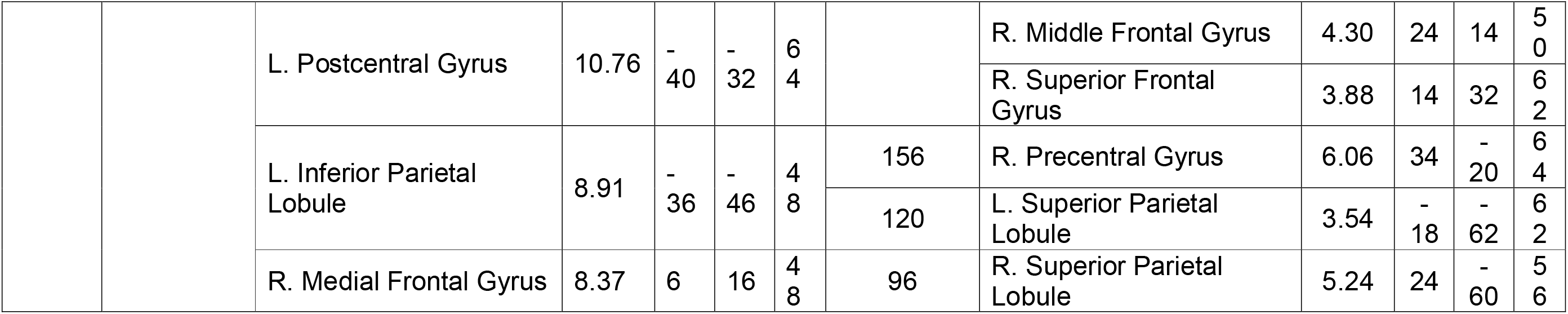
MNI coordinates of significant clusters during the five TMS conditions and the HF TMS.

### The dependency of fMRI signal changes on TMS frequency and pulse number

We further tested if the fMRI signal differed in response to changes in TMS frequency and pulse number. No significant change in the fMRI signal was found between the two conditions with high-frequency (30- and 10-Hz) TMS pulse delivery. In comparison to the LF condition, fMRI signals in the HF condition were significantly increased at the SMA, the caudal cingulate cortex, and left SMC (**Figure 4a** and **Table 2**).

**Figure 4.**
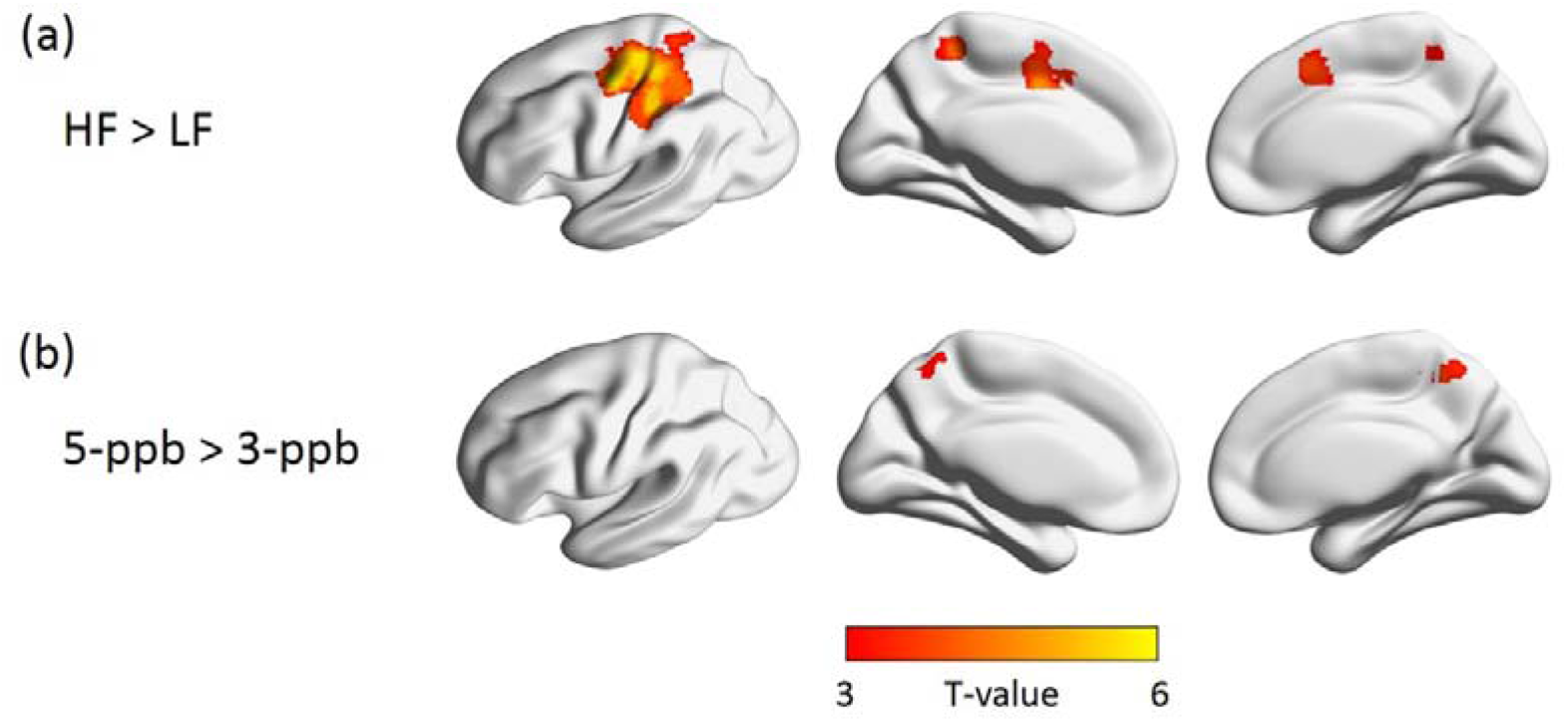
Maps for fMRI signal differences in comparisons between the HF and LF TMS conditions (a) and between 5-ppb and 3-ppb conditions (b). (a) In comparison to the LF condition, significant fMRI signal increases in the HF condition were found in the SMA and left SM. (b) A higher fMRI signal was found at the precuneus in the comparison between 5-ppb and 3-ppb TMS. The threshold was FDR-adjusted *p* < 0.05.

**Table 2.** MNI coordinates of significant clusters of the comparisons of the stimulation frequency and the pulse number.

| Effect | Cluster size | Anatomical label | Max T | X | Y | Z |
| --- | --- | --- | --- | --- | --- | --- |
| HF > LF | 2461 | L. Precentral Gyrus | 6.09 | -34 | -24 | 56 |
|  |  | L. Precentral Gyrus | 5.91 | -38 | -10 | 58 |
|  |  | L. Postcentral Gyrus | 5.58 | -40 | -32 | 64 |
|  |  | L. Superior Frontal Gyrus | 4.43 | -32 | -12 | 68 |
|  |  | L. Inferior Parietal Lobule | 4.23 | -36 | -46 | 48 |
|  |  | L. Precuneus | 4.10 | -10 | -42 | 56 |
|  | 813 | L. Cingulate Gyrus | 5.70 | 0 | -2 | 44 |
|  |  | L. Superior Frontal Gyrus | 5.29 | -8 | -4 | 68 |
|  |  | R. Medial Frontal Gyrus | 3.86 | 6 | 16 | 48 |
| 5-ppb > 3-ppb | 282 | R. Precuneus | 5.21 | 2 | -56 | 56 |
|  |  | L. Precuneus | 4.00 | -2 | -58 | 58 |

We found that 5-ppb TMS elicited significantly larger fMRI signals at the precuneus than 3-ppb TMS (**Figure 4b** and **Table 2**). To ensure that the positive fMRI response associated with the number of pulses per burst was not due to the negative fMRI in the 3-ppb condition relative to that in the 5-ppb condition, we examined the activity pattern of the precuneus across conditions. Negligible fMRI signal changes at the precuneus were found in the LF condition and the 3-ppb conditions. This signal substantially increased in the TMS conditions with 5-ppb (**Figure 5**).

**Figure 5.**
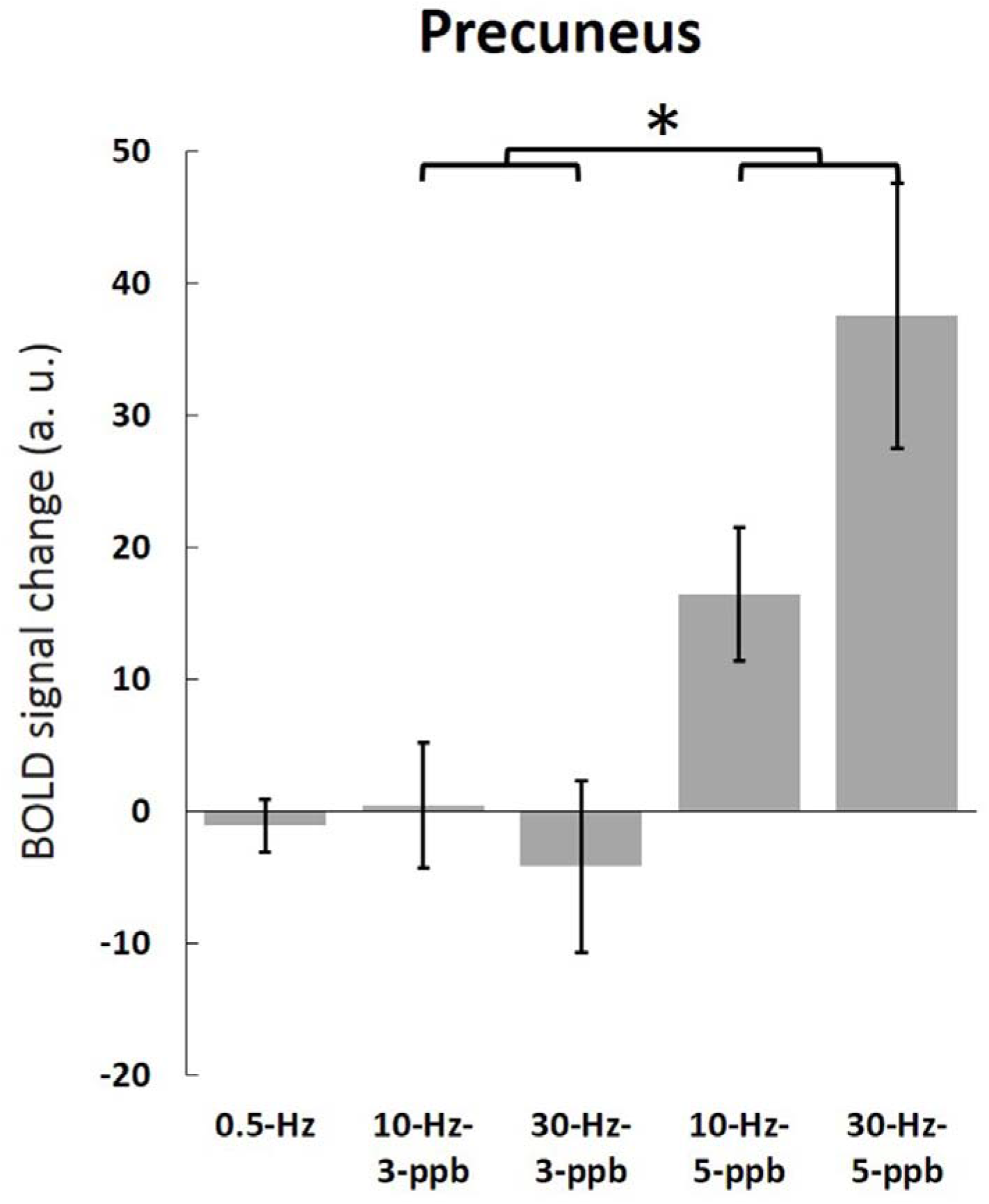
Functional MRI signal changes in TMS conditions at the precuneus. Error bars denote the SEM of fMRI signal changes in comparison to the off blocks.

### The elicited fMRI signal by TMS within the sensorimotor network

The sensorimotor network was revealed by the distribution of the significant fMRI signal changes in the finger-tapping task (**Figure 6** and **Table 3**). These areas included bilateral SMC, the SMA, bilateral insula, left supramarginal gyrus, left putamen, and cerebellum. We further examined the brain response to TMS modulations at these areas (brain areas indicated by blue contours in **Figure 3**). Because of limitations to the imaging area due to the 8-channel MRI receiver coil, we only extracted fMRI signals at the left SMC and SMA for this analysis. The average fMRI signals and mean standard errors for each ROI are shown in **Figure 7**.

**Figure 6.**
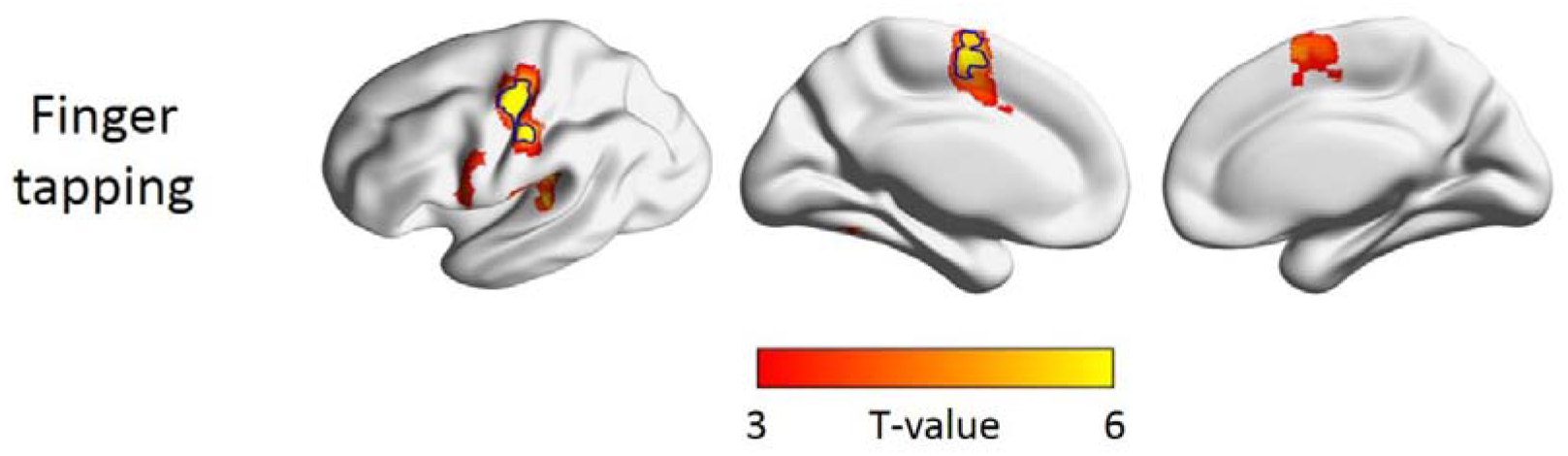
Maps of significantly increased fMRI signals in the finger-tapping task. Bilateral SMC, the SMA, bilateral insula, and left supramarginal gyrus showed significantly changed fMRI signals in the finger-tapping task. The thresholds for these maps are FDR-adjusted *p* < 0.05. Blue contours indicate the activated brain areas in the finger-tapping task.

**Table 3.** MNI coordinates of significant clusters during the finger tapping task.

| Effects | Cluster size | Anatomical label | Max T | X | Y | Z |
| --- | --- | --- | --- | --- | --- | --- |
| Finger-tapping | 1140 | R. Cerebellum | 12.77 | 20 | -54 | -24 |
|  |  | L. Cerebellum | 8.44 | -24 | -58 | -20 |
|  | 525 | L. Precentral Gyrus | 12.50 | -38 | -22 | 44 |
|  |  | L. Postcentral Gyrus | 8.86 | -36 | -38 | 52 |
|  | 631 | R. Precentral Gyrus | 12.01 | 50 | -14 | 46 |
|  | 146 | L. Supramarginal Gyrus | 6.35 | -54 | -26 | 14 |
|  | 212 | R. insula | 6.13 | 44 | 16 | -2 |
|  | 366 | L. Medial Frontal Gyrus | 6.09 | -4 | -4 | 58 |
|  |  | R. Superior Frontal Gyrus | 4.58 | 8 | 8 | 60 |
|  | 160 | L. Putamen | 5.49 | -26 | -4 | 6 |
|  |  | L. Insula | 5.18 | -44 | 0 | 4 |

**Figure 7.**
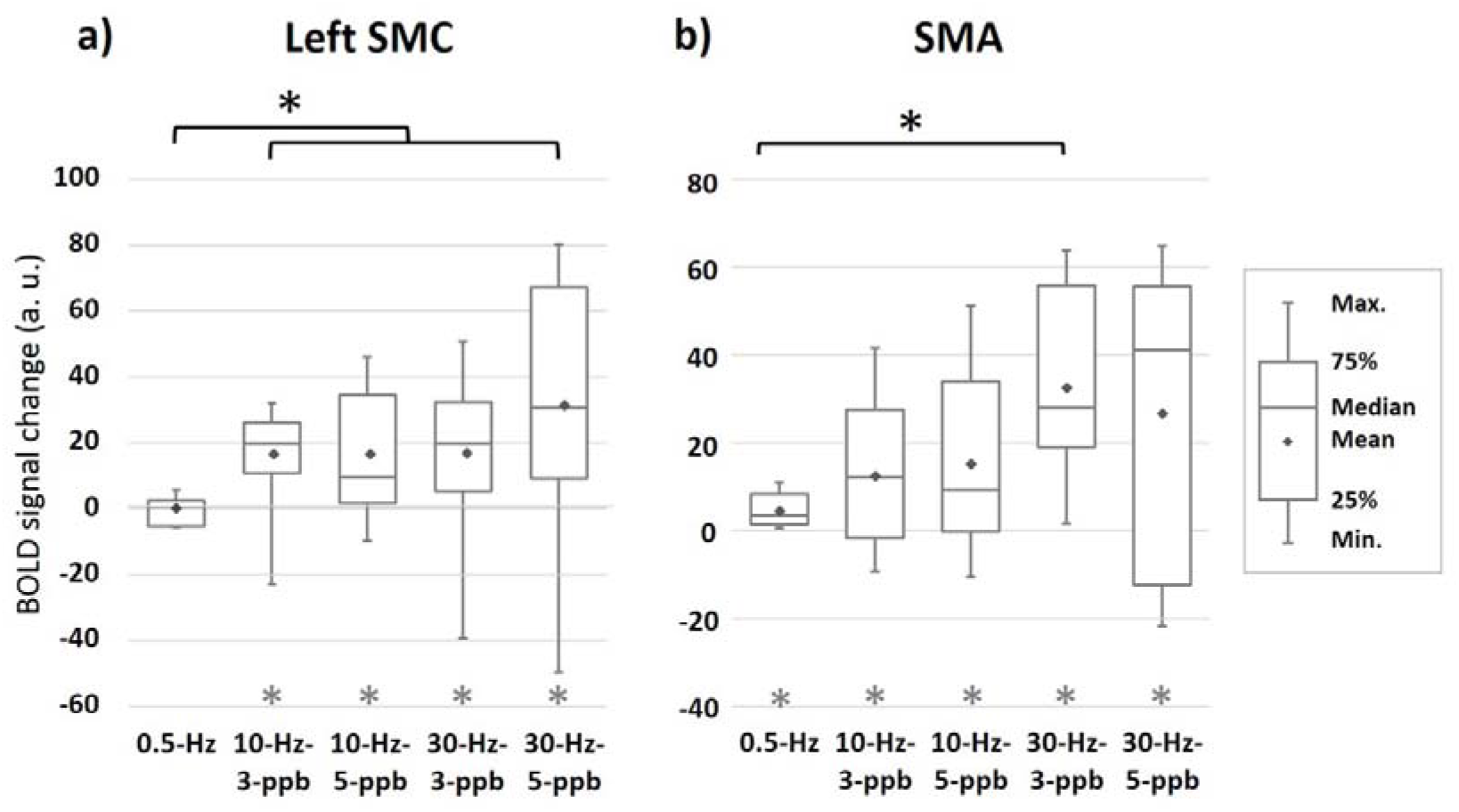
Region-of-interest analysis at the left SMC (a) and SMA (b). (a) At the left SMC, positive fMRI signals were found in each 10- and 30-Hz condition. The fMRI signal in the 0.5-Hz condition was not significantly different from that in the resting condition. The fMRI signal at left SMC in the grouped HF condition is larger than that in the LF (0.5-Hz) condition. (b) The fMRI signals at the SMA were significantly increased in all TMS conditions. Relative to the LF condition, a significantly increased fMRI signal at SMA was found in the 30-Hz–3-ppb condition relative to the 0.5-Hz condition. A gray star symbol above the horizontal axis denotes that the average fMRI signal within the region is significantly larger than that in the resting condition. A black star denotes the significant difference in the comparison between conditions indicated by the black square bracket.

The left SMC (**Figure 7a**) around the TMS target site showed significant differences in the signal variance across all five conditions (Levene’s test of variance: *F*_(4,40)_ = 3.511, *p* = 0.015). However, the variances and means of fMRI signal did not show any difference between the 10- and 30-Hz conditions (Levene’s test of variance: *F*(3,32) = 1.893, *p* = 0.151; two-way repeated-measures ANOVA: stimulation frequency: *F*_(1,8)_ = 1.296, *p* = 0.288; pulse number: *F*_(1,8)_ = 1.436, *p* = 0.265; the interaction between frequency and pulse number: *F*_(1,8)_ = 1.028, *p* = 0.340). Hence, we grouped the fMRI signals in both 10- and 30-Hz conditions as the HF condition for further analysis. Compared with the LF condition, a larger variance in the fMRI signal was found in the HF condition (*F*-test: *F*_(8,8)_ = 26.161, *p* < 0.001). The averaged fMRI signal in the HF condition was also statistically larger than that in the LF condition (Wilcoxon signed-ranks test: *Z* = −2.073, *p* = 0.038). The medians of the fMRI signal were 0.46 (−5.22 and 2.60 for the first and the third quartiles, respectively) and 21.28 (16.20 and 35.05 for the first and the third quartiles, respectively) for the LF and HF conditions, respectively. Importantly, the fMRI signal in the LF condition was not significantly different from the “off” blocks (mean ± standard deviation: −0.60 ± 4.03; two-tailed *t*-test: *t*_(8)_ = −0.446, *p* = 0.667), suggesting that the LF TMS did not lead to any detectable fMRI signal change at the SMC.

The fMRI signals at the SMA (**Figure 7b**) exhibited different variances across five conditions (Levene’s test of variance: *F*_(4,40)_ = 7.223, *p* < 0.001). Because the variances of fMRI signals were different between 10- and 30-Hz conditions (Levene’s test of variance: *F_(3,32)_* = 3.345, *p* = 0.031), we performed four separate comparisons to examine whether the variance either in a 10- or 30-Hz condition was significantly larger than that in the 0.5-Hz condition. Compared with each HF condition, we found a smaller variation in the fMRI signal at the SMA in the LF condition (*F*-test: all *p* < 0.001). The mean amplitudes of the fMRI signals at the SMA were significantly different among conditions (Friedman test: Friedman’s Q = 11.289, *p* = 0.024). Post-hoc comparisons revealed that the fMRI signal in the 30-Hz–3-ppb condition was significantly larger than that in the LF condition (Wilcoxon signed ranks test: *Z* = – 2.547, *p* = 0.011). The fMRI signal medians were 27.91 (18.94 and 55.93 for the first and the third quartiles, respectively) and 3.52 (1.44 and 8.36 for the first and the third quartiles, respectively) for the 30-Hz–3-ppb and LF TMS conditions, respectively. In all TMS conditions, the SMA showed significantly increased fMRI signal (two-tailed *t*-test: all *p* < 0.05) relative to the off blocks.

## Discussion

We systematically explored how fMRI signals change when TMS pulse sequences are designed to elicit inhibitory and excitatory modulations evidenced by MEP changes. To the best of our knowledge, this is the first study investigating fMRI signal changes in response to inhibitory and excitatory modulations by TMS. Three main findings of this study are: (1) The spatial distribution of brain activity in response to TMS intervention is comparable to that induced by voluntary finger tapping (**Figures 3** and **6**), consistently with our first hypothesis. (2) The fMRI signal change in response to inhibitory and excitatory TMS modulations is asymmetric: The excitatory HF TMS significantly increased the fMRI signal at the SMC, while the inhibitory LF TMS did not lead to detectable change (**Figures 3** and **7**). Notably, fMRI signal changes were larger at SMA than at SMC. This result contradicts with our second hypothesis, where we hypothesized that the fMRI signal changes due to TMS modulation at regions in a functional network are similar. (3) The anterodorsal precuneus, anatomically connected to the SMA [21] but outside the typical sensorimotor network, showed the pulse-number-dependent responses to TMS. We observed a significant increase in the fMRI signal between conditions of five and three TMS pulses delivered in a sub-second time scale (**Figure 4b**). Taken together, TMS activated brain areas differently based on the TMS parameters within a subsecond time scale when the dosage in the 30-s interval stayed the same.

Overall, our results can be discussed in the context of the excitatory–inhibitory balance of canonical microcircuits [22, 23]. Inhibitory neuromodulation may increase or decrease energy consumption depending on the relative contribution by interneuron spiking rates [24], the inhibitory postsynaptic potential [25], the inhibitory neuron density [26], and neurovascular coupling [27]. These factors all account for the insignificant and significant fMRI signal changes at different brain areas under TMS modulation. Further studies with concurrent electrophysiological, metabolic, and cerebral blood flow/volume measures may disentangle this complexity by building a model to predict the local brain responses to TMS.

A key finding of this study was that fMRI signal change around the stimulated M1 was only significant when TMS was delivered at a high frequency (>= 10 Hz) for eliciting the excitatory neuromodulation. The observation that excitatory TMS modulation elicited substantial increases in fMRI signal was consistent with previous PET studies, where a noticeable increase in the local glucose metabolism and regional cerebral blood flow were caused by delivering TMS pulses at a rate higher than 10 Hz [28–30]. Conversely, inhibitory TMS modulation, as measured by the suppressed MEP amplitudes, caused no significant fMRI signal change at the SMC. The inhibition of cortical excitability can be explained by the reduction of the summed synaptic activity [31]. Specifically, inhibitory synapses are fewer than excitatory ones [32–34]. Therefore, inhibitory modulation may be less likely to elicit fMRI signal changes than excitatory modulation [35]. Similar observations that inhibitory modulation fails to cause significant fMRI response at the stimulation site was also reported in other TMS–fMRI studies [6, 9, 10].

In line with our first hypothesis, we found spatial correspondence of SMC and SMA activation between TMS intervention and the finger-tapping task (**Figures 3** and **6**). However, compared to the SMC, the SMA exhibited different fMRI signal changes in response to TMS. While the SMC exhibited significant fMRI signal changes only with excitatory TMS modulation, the SMA showed significantly increased fMRI responses to both excitatory and inhibitory TMS modulations. The SMA is densely connected with M1 [36]. This structural feature supports its functional involvement in the processing of self-generated movements [37, 38] and resting-state functional connectivity [11]. The propagation of TMS from M1 to SMA has been reported in concurrent TMS and fMRI studies [6, 7, 10] and direct electric stimulation research [39] regardless of the detectable activity at M1. The connectivity-based spread of the TMS effect may be attributed to neurotransmitter release through the projections from the stimulated M1 [40].

With a gradual increase of TMS pulses delivered in a short time scale (1 s in this study), more brain areas were involved progressively. With the controlled TMS dosage over a larger time scale (30 s), we observed sequential recruitment of activated brain areas, including SMC, SMA, and precuneus. The precuneus was recently found deactivated when stimulating M1 with a train of 12 pulses at 1 Hz [41]. In our results, the anterodorsal part of the precuneus showed a significant fMRI signal increase in the 30-Hz–5-ppb condition. The fMRI signal changes at precuneus were found associated with the pulse number delivered in the sub-second range. In the HF condition, delivering pulses with 5-ppb elicited a stronger fMRI signal at the precuneus than 3-ppb. Despite not being considered as a typical area in the sensorimotor network, the anterodorsal precuneus anatomically connects to the SMA [21] and coactivates with it in the resting state [42]. Thus, we hypothesize that the precuneus activity was due to the propagation of TMS stimulation effect from the SMA, rather than from the SMC, because no direct fiber tracts were found between the precuneus and primary sensory regions [43]. The sequential recruitment of the activated brain areas from the vicinity of the stimulation site to an anatomically connected area and subsequently to an extra-network area suggests that the spatial extent of brain activation induced by local TMS depends on TMS pulse dosage within a short time scale. This result has important implications for the clinical and neuroscientific use of TMS to modulate deep brain areas distributed in a functional network by stimulating a superficial cortical area. For example, TMS has been applied to treat major depressive disorder. Studies on depressed adults have suggested that the subgenual anterior cingulate cortex (sgACC), nucleus accumbens (NAcc), and functionally connected dorsolateral prefrontal cortex (DLPFC) are pivotal for treatment efficacy [40, 44–46]. Our results suggest the use of an individualized TMS protocol for each depression patient with different TMS pulse numbers to control the propagation of brain activity from DLPFC to sgACC or NAcc. This strategy offers the opportunity to increase the efficacy of TMS treatment to depression patients and to reduce dropout rates.

In our study, a customized 8-channel MR receiver coil array was built to mount the TMS coil for precisely targeting. The customized MR receiver coil improved the sensitivity of detecting fMRI signals [16, 47]. However, the sensitive region is limited around the TMS target. This coverage limited the assessment of propagation of TMS modulation across the hemisphere. For studies aiming to investigate the distributed brain activities, a receiver coil array with a wider coverage is needed.

The current TMS setup activates a mixture of neurons with diverse neurotransmitters, distinct response profiles, and different sensitivity [48]. As fMRI signal is only secondary to neuronal activity, EEG measurements with time- and frequency-domain analyses can better characterize the brain responses to complement the information brought by fMRI signals. Taking the concurrent recording of EEG with TMS–fMRI can further disentangle the complexity of neurovascular coupling across time and spatial scales to inform how the brain responds to non-invasive excitatory and inhibitory neuromodulations locally and globally.

## Declaration of Interests

The authors declare no competing interests.

## Acknowledgments

This work was partially supported by the Academy of Finland (No. 298131), and the Natural Sciences and Engineering Research Council of Canada (RGPIN-2020-05927).

## Notes

### Competing Interest Statement

The authors have declared no competing interest.

